# External task switches activate default mode regions without enhanced processing of the surrounding scene

**DOI:** 10.1101/2024.03.04.583347

**Authors:** Ashley X Zhou, John Duncan, Daniel J Mitchell

## Abstract

Default mode network (DMN) activity, measured with fMRI, typically increases during internally directed thought, and decreases during tasks that demand externally focused attention. However, Crittenden et al. (2015) and Smith et al. (2018) reported increased DMN activity during demanding external task switches between different cognitive domains, compared to within-domain switches and task repeats. This finding is hard to reconcile with many dominant views of DMN function. Here, we aimed to replicate this DMN task-switch effect in a similar paradigm and test whether it reflects increased representation of broader context, specifically of a scene presented behind the focal task. In Core DMN, we found significant activity for all task switches, compared to task repeats, and stronger activity for switches between rest and task. Although the content of the background scene was attended, recalled, and neurally decodable, there was no evidence that this differed by switch type. Therefore, external task switches activated DMN without enhanced processing of the surrounding scene. Surprisingly, DMN activity at within-domain switches was no less than at between-domain switches. We suggest that modulation of DMN activity by task switches reflects a shift in the current cognitive model and depends on the overall complexity of that model.

## 1. Introduction

We frequently encounter scenarios where we must tackle a sequence of tasks, each requiring different cognitive resources. Such situations demand that executive control processes reconfigure mental operations to implement the appropriate task-set or procedural ‘schema’ to execute current and future goals (Monsell, 2003). An effective task model must include representations of multiple rules and goal states in working memory, while tracking and foregrounding those that are relevant in the current context. Task switching, the act of shifting attention from one task to another, is cognitively demanding, as demonstrated by behavioural switch costs whereby people’s responses are substantially slower on switch trials compared to repetitions of a task (Monsell, 2003). Accordingly, in brain imaging studies, task switching is typically associated with increased activity in regions implicated in cognitive control (Braver et al., 2003; Yeung et al., 2006), such as the multiple demand system (Assem et al., 2020; Duncan, 2010) which consists of frontoparietal brain regions that respond to cognitive challenges of many kinds.

Surprisingly, recent findings show that parts of the default mode network (DMN), although typically deactivated during difficult tasks that require external cognitive control, can also be activated by task switching (Crittenden et al., 2015; Smith et al., 2018), especially during the most demanding kind of task switching in their paradigm. Similar switch-related activation of the DMN has also been demonstrated in macaques (Arsenault et al., 2018), suggesting an evolutionarily preserved function. DMN activity at task switches is consistent with more general DMN activation at event boundaries in narrative comprehension (Speer et al., 2007) and hierarchical task sequences (Wen, Duncan, et al., 2020), yet is difficult to reconcile with many prominent views of DMN function that emphasise involvement in internally-oriented cognition (Buckner et al., 2008), such as self-related thought (Andrews-Hanna et al., 2010; Davey et al., 2016; Gusnard et al., 2001), autobiographical memory retrieval and mental imagery (Addis et al., 2007; Buckner & Carroll, 2007; Hassabis & Maguire, 2009; Schacter et al., 2007), as well as social cognition (Mars et al., 2012). The difficult, externally-driven switches between fast-paced decisions on simple stimuli, that can activate the DMN (Crittenden et al., 2015; Kurtin et al., 2023; Smith et al., 2018), seemingly afford little opportunity, let alone requirement, for such modes of thought.

So, why might some task switches activate the DMN? Given a role in episodic recall (Buckner & Carroll, 2007) and memory-guided cognition (Murphy et al., 2019), one suggestion was that DMN activation might be associated with retrieving the rules of the new task following a switch; however, DMN activity does not seem to be driven by the number of rules to be retrieved (Smith et al., 2019). Here we consider a set of alternative hypotheses that all generally concern a broadening of attention (Buckner et al., 2008). As well as focusing on the immediate task at hand, passive overseeing of the environment may be critical to determine the situational context, which in turn could determine how we execute the tasks or change our priorities. An early proposal, the “sentinel hypothesis”, highlighted a role of DMN in broadly monitoring the peripheral and internal environment (Gilbert et al., 2007; Gusnard & Raichle, 2001; Hahn et al., 2007; Raichle et al., 2001; Shulman et al., 1997), hence explaining the deactivation of DMN in tasks where focused attention to foveal stimuli is prioritised. Plausibly, this could explain DMN activation at task switches: in an evolutionary sense, it would be adaptive to monitor for new threats or opportunities when encountering a change in context or initiating a new behaviour.

More recently, components of the DMN have been associated with processing ‘situation models’ that represent the overarching setting within which behaviour occurs, and associated action schemata that the setting affords (Ranganath and Ritchey, 2012). Such situation models can help to predict and constrain which actions will be appropriate, and, as with the sentinel hypothesis, such assessment of broader context may be especially beneficial when switching to a new behaviour. Thirdly, DMN activation has been frequently linked to off-task thought, such as ‘mind-wandering’(Christoff et al., 2009; Mason et al., 2007) and ‘exploratory states’ (Shulman et al., 1997). This could also be consistent with DMN increases during task switches, if a break in routine responding affords an increased likelihood of mind-wandering or other distraction from the focal task. While mind-wandering is a broad term that encompasses stimulus-independent thoughts, it is also likely to include attention to aspects of the external environment (Gilbert et al., 2007). All three hypotheses, while positing somewhat different roles for the DMN, suggest a similar prediction – that when a task switch activates the DMN this may be accompanied by a spreading of attention from the focal task to the surrounding scene.

To test this prediction, we inserted a scene image behind the focal task in a similar task switching paradigm, and assessed scene processing in four separate ways, using a combination of behavioural and neural indices: speed of response to occasional scene probes, neural representation of scene category, neural sensitivity to repeated scene exemplars, and encoding of scenes into memory, measured using a subsequent recognition test. If DMN activity at task switches reflects a broadening of attention to encompass surrounding context, then it should be accompanied by evidence of increased scene processing, relative to task repeats.

## 2. Methods

### 2.1 Participants

38 participants were recruited from the Cognition and Brain Sciences Unit’s healthy participant panel, of whom three were excluded owing to not completing the entire experiment. The 35 analysed participants were native English speakers, aged 18-45 (63% female), with normal or corrected-to-normal vision, and with no history of neurological or psychiatric disorders. Participants provided informed consent and were monetarily compensated for their time. The experiment was conducted in accordance with ethical approval granted by the Cambridge Psychology Research Ethics Committee (CPREC).

Sample size was specified in advance, based on statistical power calculations. BUCSS (Anderson et al., 2017) was used to estimate the desired sample size to replicate the key result from Crittenden et al. (2015), the main effect of switch condition (no-switch versus between-domain switch) on univariate activity in the Core DMN, which had an F value of 16.7, and sample size of 18. Using 0.05 for the alpha level of both the planned and previous study, and 0.8 for both the desired level of statistical power and the desired level of assurance, gave a minimum sample size of 21. To test for a similar effect of switch condition on background scene representation, an estimated effect size was not available based on existing literature, so we assumed a medium standardized effect size of Cohen’s d = 0.5 for a 2-tailed t-test with alpha of 0.05 and power of 0.8, yielding a minimum required sample size of 34. We recruited 38 participants on the assumption that some might need to be excluded.

### 2.2 Task Design

The experimental tasks were adapted from Crittenden *et al*. (2015) and are illustrated in Figure 1. Participants were trained before the scanning session and learned four main tasks: two from the lexical domain and two from the semantic domain, all with yes/no answers. The two lexical tasks concerned a word stimulus with one letter missing; a blue or yellow frame around the word would specify the particular judgement required, either ‘Does the letter A fit in to make a word?’ or ‘Does the letter I fit in to make a word?’. The two semantic tasks concerned a picture of an object; a red or green frame around the object would specify whether the participant should judge ‘Is it living?’ or ‘Does it fit into a shoebox?’ respectively. The overall stimulus layout is shown in Figure 1C. Each trial began with presentation of a coloured frame on one side of the screen, indicating the location of the relevant stimulus and cuing the task to be performed on it.

**Figure 1.**
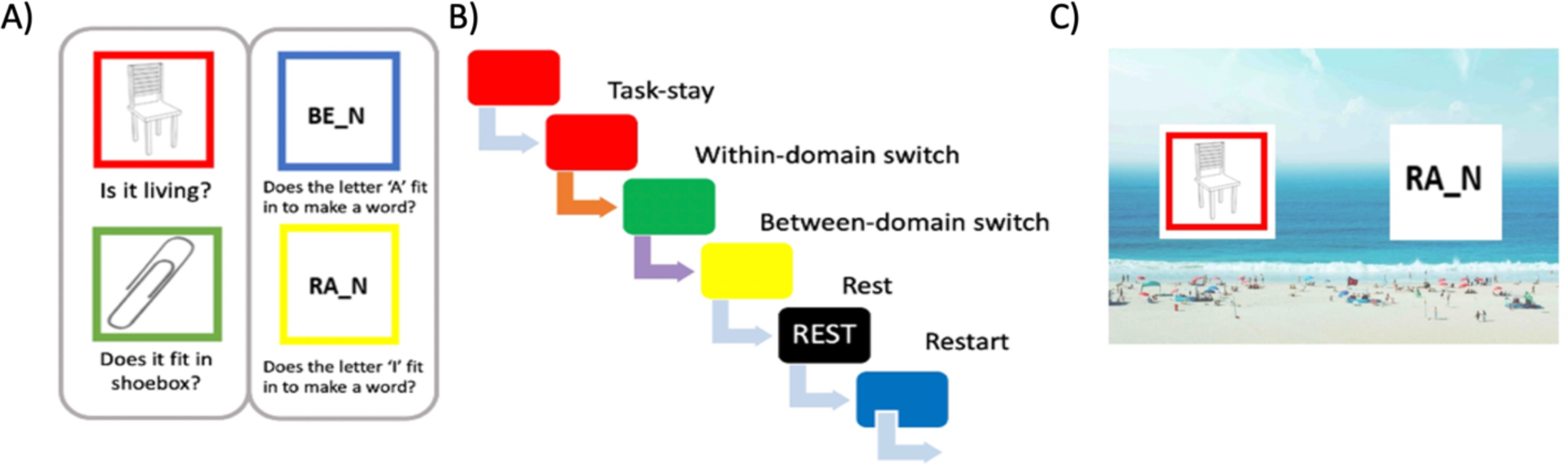
Experimental paradigm. The left panel (A) shows the four tasks and frame-colour pairs used in the experiment, organised by stimulus domain. The middle panel (B) shows an example section of a run, where the sequence of tasks creates one of each of the task-switch conditions examined in data analysis. The orange arrow indicates a within-domain switch, where the participant proceeds to a different task in the same stimulus domain; the purple arrow indicates a between-domain switch, where the participant proceeds to a task from the other stimulus domain. The right panel (C) illustrates a single trial, with two stimuli superimposed on a background scene, and the coloured frame cuing the relevant stimulus and required task.

After a delay of 0.5 seconds, a scene image (usually a beach or café; see below) was presented in the background, and 0.5 seconds after this, one word and one object stimulus were presented, one to the left and the other to the right. In the cue box was a stimulus from the category (word or object) appropriate to the cued task, while on the opposite side was a stimulus from the other category. Additional ‘rest’ trials were cued by a black frame, which indicated that the participant did not need to respond to the focal stimuli. Task stimuli were approximately 4.6 degrees of visual angle (dva) in width, and were presented at approximately 5 dva either side of fixation.

For task trials, stimuli remained on screen until a response was made by pressing one of two buttons with the index and middle fingers of their right hand; at this point, the entire display, including the background scene, disappeared. For rest trials, stimuli were presented for a duration that matched the mean duration of all preceding task trials. An inter-trial interval (ITI) of 1.5 seconds, with a central fixation cross, followed each trial.

Trials were presented in pseudorandom order, which allowed for five conditions of interest, based on transitions between successive tasks (Figure 1B). The “task-repeat” condition consisted of trials preceded by the same task and served as a baseline. The “within-domain switch” condition consisted of any trial preceded by the other task type in the same domain, and the “between-domain switch” condition consisted of trials preceded by either of the task types in the other domain. The “restart” condition consisted of all task trials that followed rest trials, which formed the “rest” condition. As restart trials involve a switch from rest to a task, we consider them as a type of “task-switch,” along with the “within-domain switch” and “between-domain switch” conditions.

In addition to the main tasks, participants were asked to identify occasional trials when the background scene contained a jungle, and to respond by pushing a third button with their left hand, instead of performing the focal task. This was to encourage constant low-level monitoring of background context, and to provide a behavioural measure of the extent of background monitoring. As mentioned above, the background image of the majority of trials contained either a beach or a café. Half of these scene images were unique to a single trial; for the remainder, each transition type was assigned a repeated image per run, which recurred on every other instance of that transition type. Image category (beach, café) remained constant between one jungle trial and the next, then, following a jungle trial, changed to the alternative category.

Participants completed four runs of 181 trials each, with tasks and scene category (beach/café) balanced across switch types (task repeat, within-domain switch, between-domain switch, restart and rest). In each run, there were 48 rest trials and 24 trials of each of the other transition types, arranged in six mini-blocks that each balanced task with transition type. There were equal numbers of tasks with either answer (yes/no), and with the frame appearing on either side of the screen, randomly ordered. The remaining trials included 18 with jungle scenes, three per mini-block and also balanced across switch types, plus unanalysed dummy trials inserted after each jungle trial and at the start of each run.

During pre-scan training, participants were shown the colours that corresponded to each rule, listened to the experimenter describe the rules, and read the instruction sheet until they understood the task. Then, participants practiced making each judgement one colour/rule pair at a time, ten times each. Participants were then asked to repeat aloud which judgement each colour cued and attempted 30 trials with intermixed cues. Finally, the experimenter explained the additional requirement to identify the occasional jungle scenes, and the participant attempted a practice block of the full task. Participants were allowed to proceed to the testing phase if their performance on the practice block exceeded 80% correct; otherwise, they attempted another block. Training lasted on average 20 minutes, after which participants were scanned for four task blocks of approximately ten minutes each.

After the fMRI scan, participants completed a computer-based memory task. For each combination of scene category and transition type (in random order per participant), participants were presented with a random arrangement of 16 scene images and were asked to identify which ones they had seen during the experiment. Four of these had appeared multiple times, four were randomly selected from the scenes that had appeared once, and eight had not been shown in the main experiment. Regardless of the accuracy of their selections, the task progressed after selecting half of the pictures. Participants were subsequently asked to sort the four repeated scene images (one per run), associated with a given switch condition, according to the run in which each was presented. Performance on the sorting task was very low and was not analysed further.

### 2.3 Behavioural data analysis

For the focal task, slow responses exceeding three standard deviations from the mean of all trials per participant were excluded. Reaction times were then averaged across correct trials per condition.

For analysis of the behavioural scene detection task only, two participants were omitted due to never responding to jungle trials during rest trials. One participant did not complete the post-fMRI memory task and so was excluded from analysis of that task.

For the subsequent scene-recognition task, accuracy was measured using the signal detection measure d’, which quantified sensitivity in discriminating scenes that had appeared in the main task (either once or multiple times) from those that had not.

### 2.4 fMRI data acquisition and pre-processing

Data were acquired on a 3T Siemens Prisma MRI scanner, fitted with a 32-channel coil. A structural T1-weighted structural scan was acquired using an MPRAGE sequence (TR 2.4s, TE 2.2ms, flip angle 8°, voxel size 0.8 × 0.8 × 0.8 mm). Task runs were scanned using T2*-weighted Echo-Planar Imaging (multiband acquisition factor 2, TR 1.2 s, TE 30 ms, flip angle 67°, voxel size 3 × 3 x 3 mm).

Pre-processing was performed in Matlab (The Mathworks Inc.) using automatic analysis (Cusack et al., 2015) to call SPM12 functions (Wellcome Department of Cognitive Neurology, London, UK). Pre-processing steps included spatial realignment of functional volumes, slice-time correction, co-registration to the T1-weighted structural, and normalization to the Montreal Neurological Institute (MNI) template brain. No smoothing was used for multivariate or ROI analyses.

### 2.5 Regions of interest (ROIs)

Primary univariate analyses focus on the Core and medial temporal lobe (MTL) subnetworks of the default mode network, where task-switch-related activity has previously been observed (Crittenden et al., 2015; Smith et al., 2018). The dorsomedial prefrontal cortex (dMPFC) subnetwork was not included, because it does not exhibit a task-switch-related response (Crittenden et al., 2015; Smith et al., 2018). ROIs for these networks were taken from Wen et al (2020), based on a subdivision of the DMN and orbital frontal–temporopolar networks from the Yeo et al (2011) 17-network parcellation, to identify voxels clusters in the vicinity of each coordinate provided in Andrews-Hanna et al. (2010). (See Wen et al. 2020 for further details.) An ROI covering the frontoparietal multiple demand (MD) network was taken from Mitchell et al. (Mitchell et al., 2016), to confirm its expected activation by task switches relative to task repeats, and activation during task relative to rest. Finally, the 17 networks from the Yeo (2011) parcellation were used as ROIs for multivariate decoding, to identify regions that carried information about scene category or novelty, averaged across conditions.

### 2.6 Univariate modelling of fMRI data

General linear models contained regressors for each combination of switch condition (task repeat, within-domain switch, between-domain switch, restart, rest), crossed with either background context (beach, café), or (separately) with background novelty (repeated, novel). Regressors were created by convolving the canonical hemodynamic response function with the duration (from scene onset to response time, capped at three standard deviations above mean) of each trial’s reaction time, for all trials per condition. Similar regressors were added to model jungle and dummy trials, which were not analysed further. Six rigid-body realignment parameters were included, along with run means, as non-convolved regressors. A temporal high-pass filter with a default cut-off period of 128 s was applied to the data and the model. These models were also used for multivariate decoding. To assess the univariate effect of task switches and rest, estimated coefficients for each switch condition were contrasted against the “task repeat” baseline condition, averaging across background scene types.

### 2.7 Multivariate decoding of fMRI data

Neural representation of background scene category (beach vs café) was examined by training a linear LDA classifier (LIBLINEAR, Fan et al., 2008) to discriminate scene category, based on activation patterns from the GLM described above, using a leave-one-run-out cross-validation approach, implemented using The Decoding Toolbox (Hebart et al., 2015). Classification accuracy was quantified as d’, separately for each Yeo network ROI. Across participants, for each ROI, the classification accuracy was first compared to chance using one-sample, 2-tailed t-tests, after averaging across all five task conditions. Significance was corrected for multiple comparisons by controlling the false discovery rate (FDR; Benjamini & Yekutieli, 2001) to be < 0.05. This served to identify a set of brain networks whose activity contained information about scene category. Classification accuracies were then averaged across these networks for each switch condition separately, which were compared using repeated measures ANOVA and paired, 2-tailed t-tests.

A similar approach was used to identify which brain networks contained information about scene novelty (averaged across task conditions), and then to compare the strength of this representation across task conditions.

Finally, given our *a priori* interest in the Core and MTL subnetworks of the DMN, and expectation that these regions would represent scene context (Smith et al., 2021), classification accuracy was also compared across conditions within the DMN subnetwork ROIs.

### 2.8 Statistical analysis strategy

For each behavioural and neural measure, we followed the same analysis strategy, to address two key questions. First, we assessed whether there was any difference between repeating a task and switching to a different task. This was tested using a paired, 2-tailed t-test that contrasted the “task repeat” condition to the average of the three switch conditions (“within domain”, “between domain”, and “restart” i.e. switch from rest to task). We expected switches to exhibit slower reaction times (Monsell, 2003), and increased activation in Core and MTL subnetworks of the DMN (Smith et al., 2018); the novel prediction was that switches would also be associated with increased background scene processing. Second, we assessed whether there was any difference between the three types of switches. This was tested using one-way repeated-measures ANOVA between the three switch conditions, to be followed by pairwise post-hoc tests if a significant effect were found. Based on Crittenden et al. (2015) and Smith et al. (2018), we expected activation in the Core and MTL subnetworks of the DMN to be lowest for within-domain switches, greater for between-domain switches, and highest for switches from rest back to task. We predicted that any differences in DMN activation might be mirrored by similar differences in background scene processing.

In addition, we ran two further tests where appropriate, to confirm the sensitivity and validity of the measures. For cases where a test versus zero was meaningful (recognition memory for scenes, and multivariate decoding of scene information), we averaged all conditions and confirmed above-chance accuracy using a one-sample, two-tailed t-test. For all measures other than main-task RT, rest was compared to the average of the four task conditions using a paired, two-tailed t-test. We expected rest to be associated with the usual activation of the DMN, and deactivation of the MD network, relative to task conditions. We also expected rest to be associated with enhanced scene processing, due to the reduction of competing focal-task demands.

To complement traditional frequentist statistical tests, Bayes factors were used to quantify evidence for the null hypothesis. Default Bayes factors were used, based on F statistics for ANOVAs (Faulkenberry & Brennan, 2023) or t statistics for t-tests (Rouder et al., 2009).

## 3. Results

### 3.1 Behavioural switch costs

For all main tasks combined, mean accuracy was 94.2%. Reaction times, averaged across correct trials, are plotted in Figure 2A for each condition. An expected behavioural switch cost was observed (Monsell, 2003), with average reaction times for combined task-switch trials significantly higher than for task-repeat trials (t_34_=5.09, p<0.01, BF_10_=4.52 x10^3^). Marginally significant differences between the task-switch conditions were observed (F_(2,68)_=3.37, p=0.04, BF_10_=0.64), although with inconclusive Bayes factor. Further paired t-tests showed that between-domain switch trials were significantly slower than within-domain (t_34_ =2.39, p=0.02, BF_10_=2.71), and restart trials (t_34_=2.91, p<0.01, BF_10_=9.12), with no significant difference between within-domain trials and restart (t_34_=0.60, p=0.56, BF_10_=0.20).

**Figure 2.**
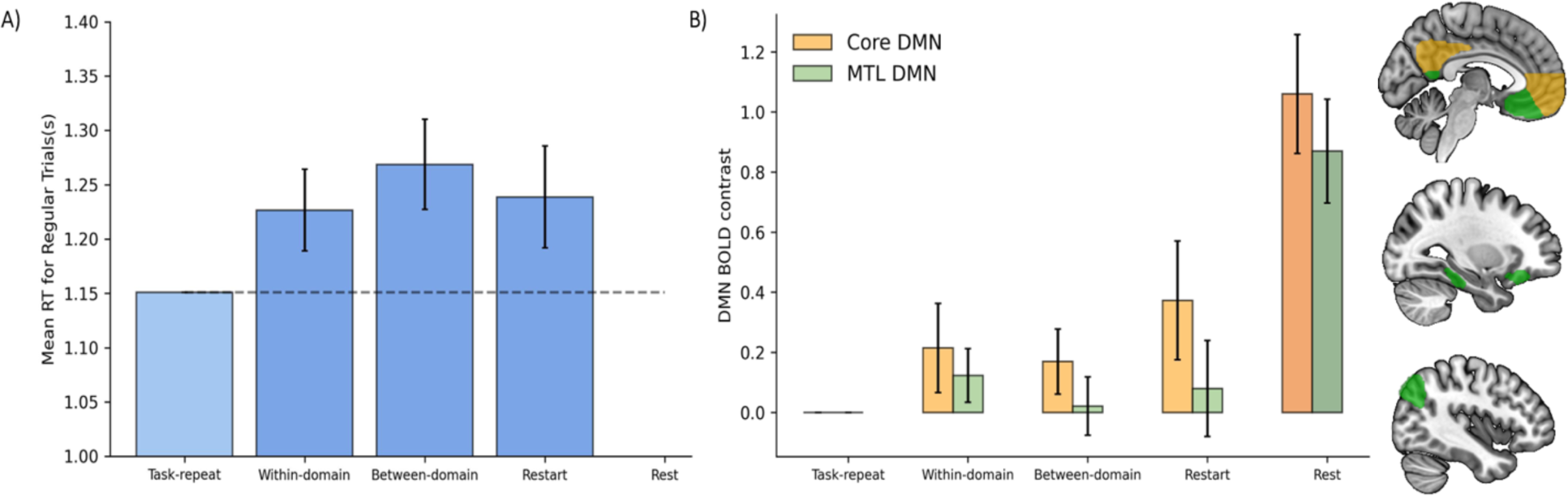
Task switches increase RT and Core DMN response, compared to task repeats. Bars in (A) show mean reaction times of correct task trials. (B) shows BOLD response in the Core and MTL DMN ROIs, for task-switch and rest trials compared to task-repeat trials. Error bars indicate within-participant 95% confidence intervals for the contrast of each condition against the task-repeat baseline.

### 3.2 Univariate DMN activity at task transitions

Given our interest in replicating the results of Smith and Crittenden (Crittenden et al., 2015; Smith et al., 2018) showing increased DMN activity for task switches compared to task repeats, we focused on the Core and MTL DMN subnetwork ROIs. The response for every condition, relative to task repeat, averaged across scene types, for both Core and MTL subnetworks is plotted in Figure 2B. An initial ANOVA, combining the three types of task transition (within-domain switches, between-domain switches and restart) and with ROI (Core, MTL) as a factor, found a significant intercept (F_(1,34)_ =9.69, p<0.01, BF=10.72), indicating that activity for task transitions was indeed higher than for task-repeat trials. There was also a significant effect of ROI (F_(1,34)_ =32.53, p<0.01, BF_10_ =7.26×10^3^)), with the relative response to task transitions being greater in the Core than MTL subnetwork. Given this effect of ROI, we proceeded with separate analyses per ROI.

Considering first the Core subnetwork, a t-test confirmed that activity for the three types of task transitions combined was higher than for task-repeat trials (t_34_=4.03, p<0.01, BF_10_=187). One-way ANOVA showed a significant difference amongst the three types of task transition (F _(2,68)_ =3.89, p=0.03, BF_10_=1.01), although the Bayes factor was inconclusive. Post hoc t-tests found no significant difference between within- and between-domain switches (t_34_=0.79, p=0.43, BF_10_=0.23), or between within-domain switches and restart trials (t_34_=1.90, p=0.06, BF_10_=1.02). However, restart trials evoked greater activity than between-domain switches (t_34_=2.35, p=0.02, BF_10_=2.52). As expected, rest trials produced higher Core DMN activity than task trials combined (t_34_=9.63, p<0.01, BF_10_=3.21 x10^9^).

For the DMN’s MTL subnetwork, there was no significant difference in response to task switches combined compared to task repeats (t_34_=1.64, p=0.11, BF_10_=0.61), although the Bayes Factor indicated only weak support for the null hypothesis. One-way ANOVA indicated no difference between the three types of task-switch conditions (F _(2,68)_ =1.16, p=0.32, BF_10_=0.09). Again, rest trials produced higher MTL activity than task trials combined (t_34_=10.56, p<0.01, BF_10_=3.82 x10^9^).

Given previous findings showing increased MD activity for task switches compared to task repeats in more traditional task-switching paradigms, we also examined activity in the MD ROI (supplementary Figure 1). In MD regions, activity for the three types of task transitions combined was indeed higher than for task-repeat trials (t_34_= 2.39, p=0.02, BF_10_=2.70). One-way ANOVA showed no significant difference amongst the three types of task transition (F _(2,68)_ =0.46, p=0.63, BF_10_=0.05). Also as expected, rest trials produced less MD activity than task trials combined (t_34_=10.87, p<0.01, BF_10_=8.59 x10^9^).

### 3.3 No effect of task transitions on behavioural measures of background scene processing

For the background-scene detection task, average accuracy was 92.7% and reaction time was 0.85 seconds (Figure 3A). Rapid responses to the occasional jungle scenes confirmed attention to the background, despite its secondary importance and irrelevance to the focal task. Reaction times for task-repeat trials did not significantly differ from trials with a transition in the focal task, averaged across transition types (t_32_=0.58, p=0.57, BF_10_=0.21). The Bayes factor indicated substantial support for the null hypothesis. An ANOVA also revealed no difference between the three transition types (F _(2,64)_=0.55, p=0.58, BF_10_=0.05), with the Bayes factor indicating strong support for the null hypothesis. A significant difference was observed between task trials and rest trials (t_32_=5.07, p<0.01, BF_10_=3.9 x10^3^), consistent with more efficient scene processing on rest trials, when competing demands of the focal task were removed. This confirms that the measure is sensitive enough to pick up behavioural differences in online scene processing.

**Figure 3.**
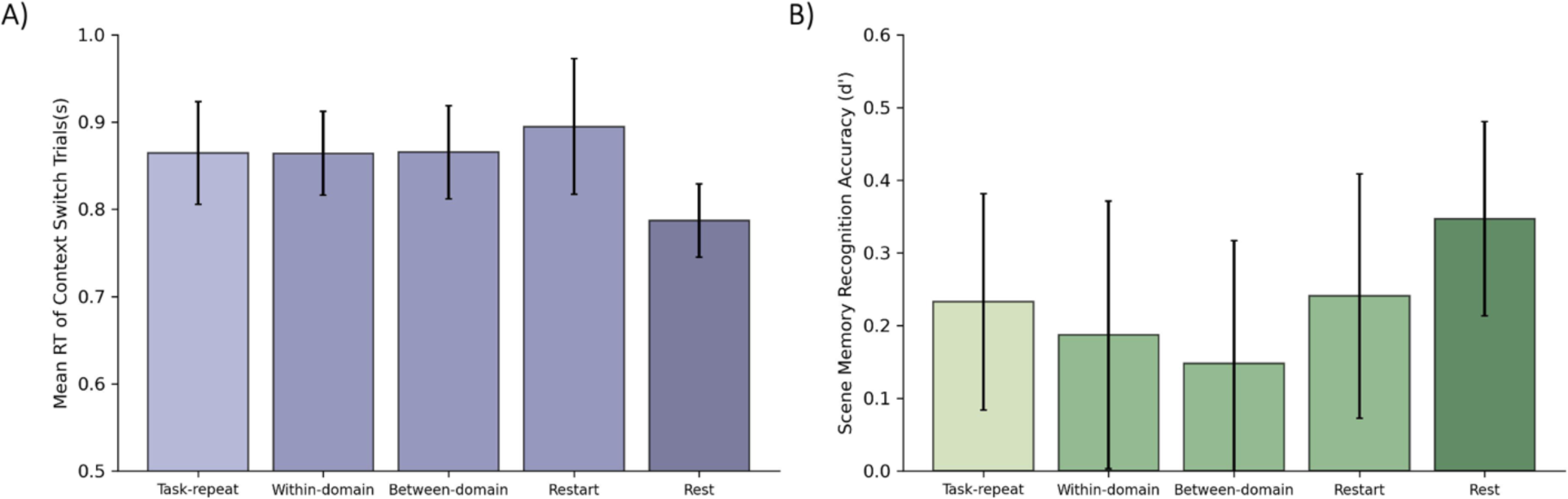
Behavioural measures of scene processing. (A) shows reaction times for accurate jungle detection trials and (B) shows scene recognition accuracy (d’) in the subsequent memory test, for scenes occurring in each task condition. Error bars indicate between-participant 95% confidence intervals.

In the post-scan memory test, averaging across all five conditions, participants recognized scenes that had been presented in the main experiment, from amongst non-presented scenes, with accuracy significantly above chance (t_33_=5.74, p<0.01, BF_10_=3.11 x10^4^); this confirmed encoding of the scenes into memory, and sensitivity of the subsequent recognition measure. Crucially, however, memory performance averaged across trials with focal task transitions showed no difference compared to task-repeat trials (t_33_=0.48, p=0.63, BF_10_=0.19). The Bayes factor indicated substantial support for the null hypothesis. There was also no difference between the three types of task transition (F _(2,66)_=0.33, p=0.72, BF_10_= 0.04), with the Bayes factor indicating strong support for the null hypothesis. A t-test showed that recognition of scenes presented on rest trials was numerically higher than for scenes presented on task trials, but with equivocal statistical significance (t_33_=2.08, p=0.05, BF_10_=1.46).

### 3.4 Task transitions did not increase MVPA measures of background scene representation

Multivariate analyses were performed to identify where in the brain the category and novelty of background scenes was decodable and therefore represented in the response pattern. We reasoned that if the DMN’s role during task switches involves representing surrounding contextual information, or more generally reflects a broadening of attention away from the focal task, then scene representation would strengthen at task switches compared to task repeat trials. Neural discrimination of scene category (beach vs café) and scene novelty were tested using a linear classifier and leave-one-run-out cross-validation. For each of the Yeo 17 Networks, the average classification accuracy was first averaged across all main task conditions and tested against zero to identify regions carrying scene information. Significant decoding of scene category (p<0.05, before FDR correction) was found in seven networks (Figure 4A) -- Visual Peripheral, Visual Central, DMN MTL, Dorsal Attention A, Control C, Dorsal Attention B, Limbic B—of which the first five survived FDR correction, confirming that scene information was represented strongly in the brain.

**Figure 4.**
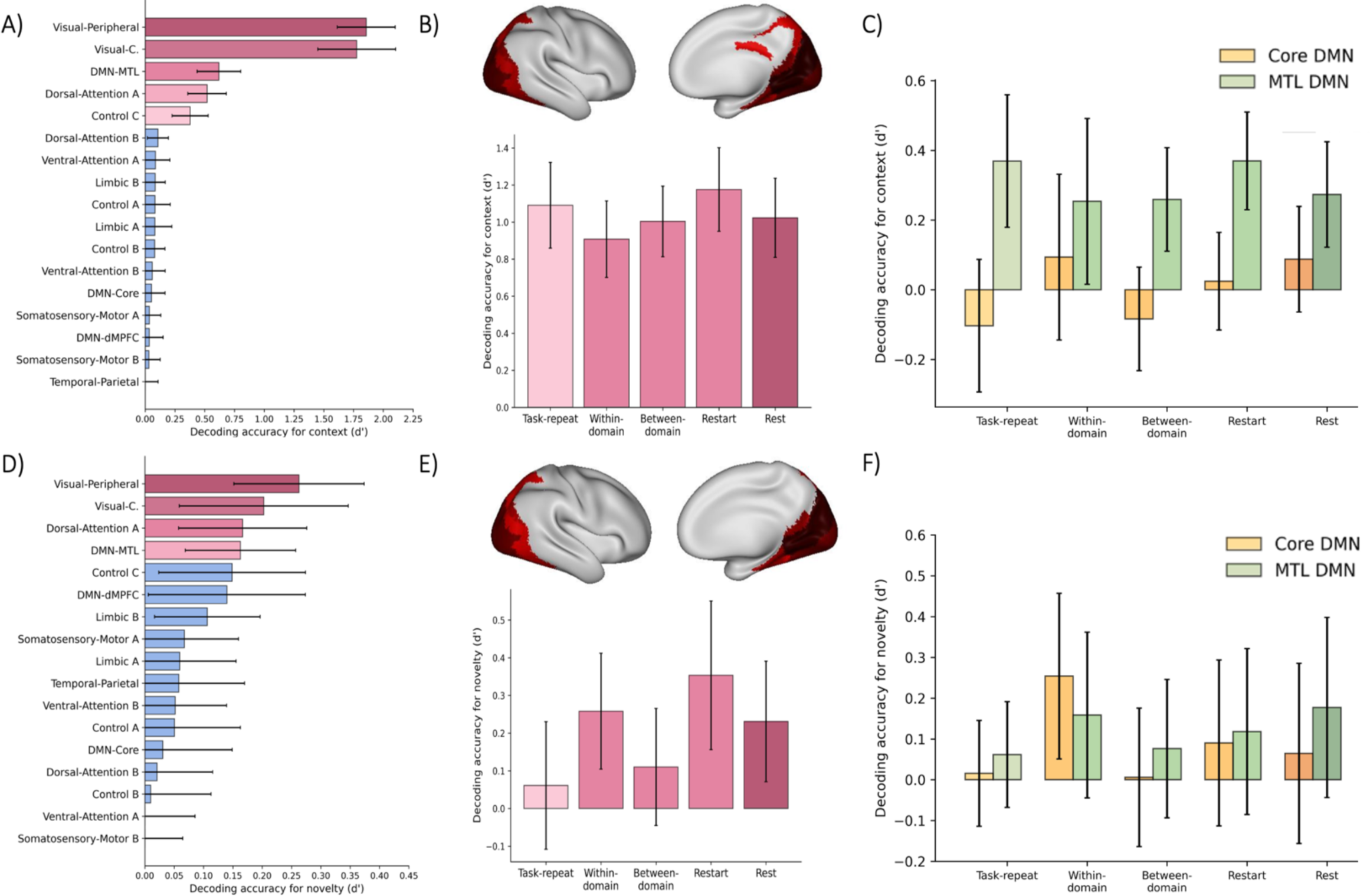
Multivariate analysis of scene category and scene novelty. The top row presents multivariate analysis of scene category (A-C). In (A), classification accuracies, averaged across conditions, are shown for each of Yeo’s 17 networks, with shades of pink indicating significant networks after FDR correction. (B) shows classification accuracy per switch condition, averaging across networks that are highlighted pink in panel A, and rendered on the right hemisphere cortical surface. Classification accuracy in MTL and Core DMN ROIs are shown in (C). The bottom row (D-F) presents multivariate analysis of scene novelty. (D) illustrates classification accuracies, averaged across conditions, for each of Yeo’s 17 networks, highlighting significant networks, after FDR correction, in shades of pink. (E) presents classification accuracy per switch condition, averaged across networks that are highlighted pink in panel D and rendered on the right hemisphere cortical surface. Classification accuracy in MTL and Core DMN ROIs are shown in (F). Error bars represent 95% between-participant confidence intervals.

Networks with significantly above-chance classification accuracies (p<0.05 after FDR correction) were then averaged, separately for each main task condition (Figure 4B). Crucially, for these averaged data, scene-category classification accuracy combined across task-transition trials did not significantly differ from task-repeat trials (t_34_=1.25, p=0.22, BF_10_=0.37). An ANOVA also indicated that scene category classification accuracy did not differ between the three types of task transition (F _(2,68)_=0.67, p=0.52, BF_10_=0.06), with the Bayes factor indicating strong support for the null hypothesis. Finally, a t-test showed no significant difference in classification accuracy between task and rest trials (t_34_=1.17, p=0.25, BF_10_=0.34), showing that scene representation during tasks was not significantly less than in rest trials.

Since we expected that DMN regions would represent scene context (Smith et al. 2021), and that this representation might strengthen along with univariate activation at a task switch, we also repeated the analysis within the *a priori* Core DMN and MTL subnetwork ROIs (Figure 4C). A two way ANOVA with factors of ROI (Core, MTL) and condition (task repeat, average of task transitions) found a significant effect of ROI, with stronger scene representation in MTL regions (F_(1,34)_ =44.05, p<0.01, BF_10_=8.63×10^4^), but no effect of condition (F_(1,34)_ =0.06, p=0.82, BF_10_=0.23), and no interaction (F_(1,34)_ =1.59, p=0.22, BF_10_=0.45). Comparing the three types of transition, again there was no main effect of condition (F _(2,68)_ =1.52, p=0.23, BF_10_=0.12), and no interaction with ROI (F _(2,68)_ =1.59, p=0.21, BF_10_=0.13). Finally, comparing rest to all task conditions, we again found no main effect of condition (F _(1,34)_ =0.21, p=0.60, BF_10_=0.25), and no interaction with ROI (F _(1,34)_ =2.31, p=0.14, BF_10_=0.61).

Next, we assessed representation of scene novelty, as an index of episodic memory processing. Neural discrimination of scene novelty was weaker than scene category, but seven networks yielded above-chance significance (p<0.05, before FDR correction) in classification accuracy (Visual Peripheral, Visual Central, Dorsal Attention A, DMN MTL, Control C, DMN dmPFC, Limbic B, of which the first four survived FDR correction), confirming that information indicating scene familiarity was retrieved from memory as scenes were processed (Figure 4D). As before, networks with significantly above-chance classification accuracies (FDR-corrected) were then averaged, separately for each main task condition (Figure 4E). Importantly, scene novelty classification combined across task-transition trials did not significantly differ from task-repeat trials (t_34_=1.52, p=0.14, BF_10_=0.54), although in this case the Bayes Factor only indicated weak support for the null hypothesis. An ANOVA also found no difference in scene novelty classification between the three types of task transition (F _(2,68)_=1.07, p=0.35, BF_10_=0.08), with the Bayes factor indicating strong support for the null hypothesis. Finally, there was a marginally significant difference in classification of scene novelty between all task trials and rest trials (t_34_=2.15, p=0.04, BF_10_=1.67), with stronger representation of scene novelty during rest.

As before, we repeated the analysis within the *a priori* Core DMN and MTL subnetwork ROIs (Figure 4F). A two-way ANOVA with factors of ROI (Core, MTL) and condition (task repeat, average of task transitions) found no effect of ROI, (F _(1,34)_=0.13, p=0.72, BF_10_=0.23), no effect of condition (F _(1,34)_=0.94, p=0.34, BF_10_=0.34), and no interaction (F _(1,34)_=0.10, p=0.76, BF_10_=0.23). Comparing the three types of transition, again there was no main effect of condition (F_(2,68)_=1.31, p=0.28, BF_10_=0.10), and no interaction with ROI (F_(2,68)_=0.70, p=0.50, BF_10_=0.06). Finally, comparing rest to all task conditions, there was no main effect of condition (F_(1,34)_=0.12, p=0.73, BF_10_=0.23), and no interaction with ROI (F_(1,34)_=0.43, p=0.51, BF_10_=0.27).

## 4. Discussion

### 4.1 Attentional broadening does not explain enhanced DMN activation at task switches

Switching from one task to another is cognitively demanding, as demonstrated by performance costs (Monsell, 2003), and is typically associated with activation of brain networks involved in executive control (Braver et al., 2003). Therefore, it is surprising that the Core subnetwork of the default mode network (DMN), which is typically deactivated during difficult external tasks, can also be activated by difficult task switches under some circumstances (Crittenden et al., 2015; Kurtin et al., 2023; Smith et al., 2018). Here we confirmed that, when people must switch between many tasks, or from a brief rest back to a task, DMN regions are activated by task switches relative to task repeats. We hypothesised that this might reflect a transient reallocation of attention at task switches, away from a narrow task focus, to encompass the broader environmental setting.

This hypothesis was tested by inserting a scene image behind the focal task and comparing behavioural and neural measures of scene processing across task conditions. Rapid behavioural responses to occasional target jungle scenes confirmed attention to the scenes. Neural decoding of scene category further confirmed that scene gist was represented strongly in the brain, while neural decoding of exemplar novelty and above-chance performance on a subsequent memory test demonstrated that details of individual scenes were encoded into memory. Despite this robust scene processing, and despite focal task switches significantly modulating both Core DMN activity and reaction times to the focal task, we found a convincing lack of evidence that task switches affected scene processing. Although this is a null result, it is supported by Bayes factors favouring the null hypothesis, and its consistency across four different neural and behavioural measures is persuasive. Thus, although the DMN has long been proposed to be involved in broadly monitoring the environment (Buckner et al., 2008; Gilbert et al., 2007; Hahn et al., 2007; Raichle et al., 2001; Raichle & Gusnard, 2001; Shulman et al., 1997) and representing situational context (Ranganath and Ritchey, 2012), and is particularly engaged when life-like scene context dictates an appropriate response (Smith et al., 2021), this role of the DMN does not seem to explain the DMN’s enhanced activity at switches between tasks.

### 4.2 Dependence of the DMN task-switch effect on overall task-set complexity

An unexpected aspect of the current results may yield insights into the requisites for recruitment of DMN during task switches. The original studies reporting DMN activity at task switches found the DMN response to be greater for switches between stimulus domains, compared with within-domain switches (Crittenden et al., 2015; Smith et al., 2018). Here, in contrast, we observed significant and comparable DMN activation for both types of switches (and for switches from rest back to task), compared to task repeats. While the reason for this difference will require further investigation, we offer one proposal. In the current study, we decreased the number of primary tasks from six (across three domains), as in the previous paradigms, to four (across two domains). This change in the overall complexity of the entire set of tasks could have affected which switches activate the DMN. Speculatively, a set of four tasks may necessitate that the tasks be mentally separated, and they may be separate enough from each other not to differentiate ‘within-domain’ from ‘between-domain’ task-switches. With six tasks, however, overloading of working memory resources might encourage a hierarchical task representation, with related task pairs grouped, and activated, together. This may explain why more traditional task switching paradigms with only two tasks do not see an increase in DMN activity at task switches at all, if two tasks can be co-activated as a single chunk and do not need to be mentally segregated. Future studies should continue to investigate the boundary conditions of the DMN task-switch effect and could specifically explore a role of task-set complexity.

A second difference from previous findings is that the DMN task-switch effect was not significant in the MTL subnetwork, but only in the Core subnetwork. Again, one might speculate that this could derive from the reduction in overall task-set complexity. Increased cognitive demands lead to expanded recruitment of regions within the MD network (Shashidhara et al., 2019), and it is possible that a similar principle could apply to DMN recruitment in the context of task switches.

### 4.3 Remaining questions regarding the role of DMN activation at task switches

Overall, the potential role of DMN in task-switches is yet to be resolved. Functions related to social processing, autobiographical retrieval and other self-focused cognition (Andrews-Hanna et al., 2010) are unlikely to be relevant in the current task paradigm. “Task-negative” perspectives on DMN function are also inadequate given the active and demanding nature of the task switches. A previous study argued against a role in retrieving the content of a new task rule (Smith et al., 2021), and the current results argue against explanations based on environmental monitoring, and other processes that reallocate attention from the focal task to the surrounding context.

Having eliminated several plausible DMN functions that may be evoked by task switches, it remains possible that the effect is more directly related to the switching process itself. For example, it could be that DMN is involved in deactivating the cognitive set for the previous task and/or activating the cognitive set required for the next task, though as discussed above, we still lack understanding of the role of overall task complexity. Involvement in cognitive switching is consistent with observations that the DMN can be activated at event boundaries in narrative comprehension (Speer et al., 2007) and hierarchical task sequences (Wen, Duncan, et al., 2020) and with DMN suppression during typical simple tasks that enforce maintenance of a single task-set. Further studies could usefully investigate whether DMN activation at task switches can be explained by a role in temporarily deactivating the preceding cognitive set. It is also unclear whether DMN is affected by, or involved in executing, a cognitive switch. Either way, the link between task switches and DMN activity is consistent with suggestions that anomalous DMN function is associated with clinical conditions characterised by dysregulated attentional switches (Fassbender et al., 2009; Sudre et al., 2017).

To conclude, we found robust evidence of background scene processing in brain and behaviour, and an increased response within the Core DMN subnetwork for behaviourally demanding switches in the foreground task. However, contrary to expectations from several prominent accounts of DMN function, we found no evidence that background scene processing was also enhanced when the foreground task changed. Therefore, the DMN response to demanding focal task switches is not readily explained by standard views of DMN function. Perhaps, instead, typical functions ascribed to the DMN, from mental simulation to environmental monitoring or mind-wandering, just like switching within a complex task set, are linked by a common denominator of involving substantial shifts within a complex cognitive model.

## Acknowledgements

Ashley Zhou, Daniel J. Mitchell and John Duncan were supported by Medical Research Council Intramural Program MC_UU_00030/7. Ashley Zhou was supported by a Gates Cambridge Trust Scholarship. The authors declare no competing financial interests.

For the purpose of open access, the author has applied a Creative Commons Attribution (CCBY) license to any Author Accepted Manuscript version arising from this submission.

---

**Supplementary figure 1.**
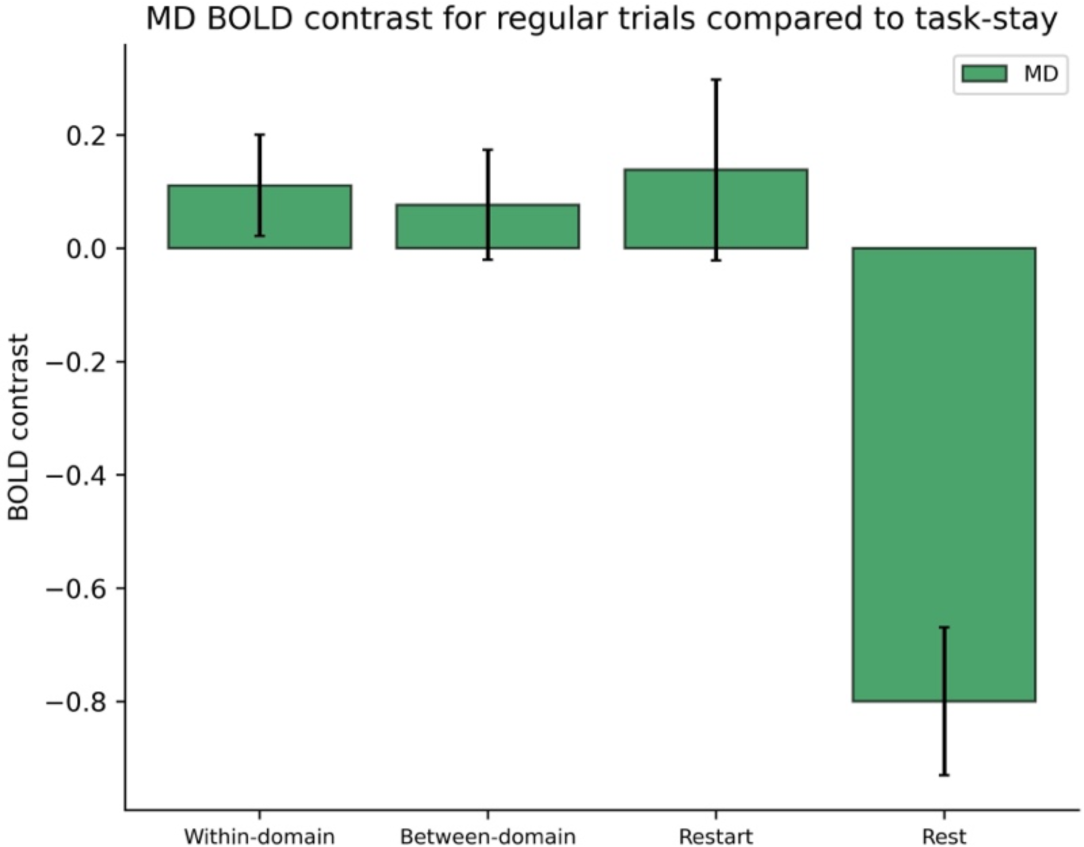
BOLD response in MD ROIs, for task-switch and rest trials compared to task-repeat trials. Error bars indicate within-participant 95% confidence intervals for the contrast of each condition against the task-repeat baseline.

## Notes

### Competing Interest Statement

The authors have declared no competing interest.

